# Mass2SMILES: deep learning based fast prediction of structures and functional groups directly from high-resolution MS/MS spectra

**DOI:** 10.1101/2023.07.06.547963

**Authors:** David Elser, Florian Huber, Emmanuel Gaquerel

**Affiliations:** Institut de Biologie Moléculaire des Plantes du CNRS, Université de Strasbourg; University of Applied Sciences Düsseldorf

**Keywords:** Structure prediction, MS/MS spectra, functional groups, deep learning, SMILES

## Abstract

Modern mass spectrometry-based metabolomics generates vast amounts of mass spectral data as part of the chemical inventory of biospecimens. Annotation of the resulting MS/MS spectra remains a challenging task that mostly relies on database interrogations, *in silico* prediction and interpretation of diagnostic fragmentation schemes and/or expert knowledge-based manual interpretations. A key limitation is additionally that these approaches typically leave a vast proportion of the (bio)chemical space unannotated. Here we report a deep neural network method to predict chemical structures solely from high-resolution MS/MS spectra. This novel approach initially relies on the encoding of SMILES strings from chemical structures using a continuous chemical descriptor space that had been previously implemented for molecule design. The deep neural network was trained on 83,358 natural product-derived MS/MS spectra of the GNPS library and of the NIST HRMS database with addition of the calculated neutral losses for those spectra. After this training and parameter optimization phase, the deep neural network approach was then used to predict structures from MS/MS spectra not included in the training data-set. Our current version, implemented in the Python programming language, accurately predicted 7 structures from 744 validation structures and the following 14 structures had a *Tanimoto* similarity score above 0.9 when compared to the true structure. It was also able to correctly identify two structures from the CASMI 2022 international contest. On average the *Tanimoto* similarity is of 0.40 for data of the CASMI 2022 international contest and of 0.39 for the validation data-set. Finally, our deep neural network is also able to predict the number of 60 functional groups as well as the molecular formula of chemical structures and adduct type for the analyzed MS/MS spectra. Importantly, this deep neural network approach is extremely fast, in comparison to currently available methods, making it suitable to predict on regular computers structures for all substances within large metabolomics datasets.

## Introduction

One of the major challenges in current metabolomics experiments is the illumination of the so called dark matter (“unknown unknowns”), which currently corresponds to the largest proportion of data analysis results, even with state-of-the-art computational methods (Beniddir et al., 2021; Aksenov et al., 2017; da Silva et al., 2015). The standard approach to retrieve high quality annotations is by spectral library matching which generally uses cosine similarity, but also alternative metrics have been developed recently, such as Spec2Vec (Huber et al., 2021a), MS2deepscore (Huber et al., 2021b) or SIMILE (Treen et al., 2022). Due to the limited number of entries in authentic standard-based spectral libraries, *in silico* fragmentation approaches have emerged such as Metfrag (Ruttkies et al., 2016), CFM-ID (Wang et al., 2021), MassFormer (Young et al., 2021) or QCxMS (Koopman and Grimme, 2021), in order to mine chemical structure libraries. Substructural information on unknown molecules can further be retrieved up to a limited extent by programs such as MESSAR (Liu et al., 2020) or MS2LDA (Wandy et al., 2018). Other approaches such as CSI:FingerID (Dührkop et al., 2015), MIST (Goldman et al., 2022) or DeepEI (Ji et al., 2020) use the generation of fingerprints from spectra to retrieve annotations from chemical structural databases. In terms of MS/MS data classification, fingerprints and similarity metrics can be used to create molecular networks as pioneered by Global Natural Products Social molecular networking (GNPS) (Wang et al., 2016), which may further give insights into main metabolic classes present within a dataset. Metrics used for molecular networking are also central to retrieve annotations by database searches. Finally, it is now possible to retrieve hierarchically-organized class-based annotations which are based on the ClassyFire (Djoumbou Feunang et al., 2016) chemical ontology with the use of the deep neural network classifier CANOPUS (Dührkop et al., 2021).

The computational prediction of molecular structures solely from mass spectra has long been envisioned as a Holy Grail in mass spectrometry, with first attempts to use artificial intelligence dating back to the launch of the DENDRAL project in 1965 (Buchanan and Feigenbaum, 1978). A seemingly logical approach to structure prediction would be to calculate the molecular formula of a molecule using SIRIUS (Dührkop et al., 2019) or BUDDY (Xing et al., 2022) and then generate all possible structures with structure generators such as MAYGEN (Yirik et al., 2021) or MOLGEN (Kerber et al., 2005), but this rapidly translates into a combinatorial explosion even for relatively small molecules. A way to circumvent this bottleneck and avoid the generation of all possible molecules is to use a continuous chemical descriptor space such as developed by Gómez-Bombarelli et al. (2018) and Winter et al. (2019). Two tools have recently emerged for *de novo* structure prediction from high resolution MS/MS spectra, MSNovelist (Stravs et al., 2022) and Spec2mol (Litsa et al., 2021), While the latter makes use of such a continuous descriptor space approach as above described. Spec2mol uses a 1-D convolutional neural network to create a latent representation of the spectra and a encoder-decoder architecture with gated recurrent units (GRU) trained on a translation task from random to canonical SMILES as employed by Winter et al. (2019). MSNovelist uses the fingerprint representation and molecular formula obtained by SIRIUS to feed a recurrent neural network that will then predict a set of SMILES which are then further ranked by scoring their probability. Another recently published tool is MS2prop, which applies transformer like architectures to predict 10 chemical properties of unknowns with high accuracy (Voronov et al., 2022a).

Despite having been reported as part of peer-reviewed articles, codes to Spec2mol and MS2prop are not available for the general public and publicly released MSNovelist relies on SIRIUS which makes it computationally demanding when processing large datasets because as it still relies on hand crafted heuristics and kernel functions. Finally, none of the available tools can predict a set of functional groups predicted to be present in a given molecule, even though it is known that direct structure prediction alone is prone to errors. Here, we report an open-sourced deep learning model that is able to quickly predict structures as SMILES strings, the presence of 60 functional groups, the adduct type as well as to give an estimation of the number of different atoms in a given molecule.

## Methods

Spectral libraries were downloaded (16.12.2022) from GNPS (Wang et al., 2016) and the NIST 2020 HRMS database was purchased by the Institute of Molecular Biology of Plants, CNRS | University of Strasbourg (IBMP). The NIST database was preprocessed with a script from MassFormer (Young et al., 2021) to correct corrupted .mol files. For the NIST and the BMDMS-NP (Lee et al., 2020) (which is contained within GNPS) the composite spectra were calculated to account for the acquisition of these spectral at several collision energies (see also **Data and code availability**). Early access to mFam Consortium Staging Database was kindly provided by Chimmiri Anusha, Steffen Neumann and Gerd Balcke. The training data was preprocessed with matchms (version 0.11.0) package (Huber et al., 2020) and rdkit (Landrum, 2010). Spectra from low-resolution mass spectrometers were excluded. Noise signals were discarded by reducing the number of peaks to 250 and corresponding neutral losses were calculated, resulting in a maximum of 500 peaks (see also **Data and code availability**). Only positive ion mode spectra with single charges were considered for training, also less frequent adducts were discarded first in pandas (https://pandas.pydata.org/) and then manually inspected and harmonized in OpenRefine (3.5.0). For validation, 744 unique spectra were randomly selected based on Inchikey and all Inchikey corresponding spectra were then discarded from the training dataset, resulting in a final training dataset of 83,358 spectra with 18 different adducts (**Table 1**).

**Table 1.** Adducts included in the Mass2SMILES training data, with their delta to the actual mass of the molecule. The encoded numbers depicted here can be used to translate the predicted adducts.

| Adduct | Delta | Encoding |
| --- | --- | --- |
| $[M+H-C_2H_4O_2]^+$ | -60 | 0 |
| $[M-3H_2O+H]^+$ | -54 | 1 |
| $[M-2H_2O+H]^+$ | -36 | 2 |
| $[M-H_2O+H]^+$ | -18 | 3 |
| $[M-NH_3+H]^+$ | -17 | 4 |
| $[M]^+$ | | 5 |
| $[M+H]^+$ | +1 | 6 |
| $[M+H+2i]^+$ | +1 + Isotopes | 7 |
| $[M+NH_3]^+$ | +17 | 8 |
| $[M+NH_4]^+$ | +18 | 9 |
| $[M+Na]^+$ | +23 | 10 |
| $[M+H+CH_3OH]^+$ | +33 | 11 |
| $[M+K]^+$ | +39 | 12 |
| $[2M+H]^+$ | $2x + 1$ | 13 |
| $[2M+H+2i]^+$ | $2x + 1$ + Isotopes | 14 |
| $[2M+NH_4]^+$ | $2x + 18$ | 15 |
| $[M-H+2Na]^+$ | +46 | 16 |
| $[2M+Na]^+$ | $2x + 23$ | 17 |
| $[2M+K]^+$ | $2x + 39$ | 18 |

Spectra were encoded by sinusoidal encodings inspired by Voronov et al. (2022b) with 256 dimensions and a precision of two decimals. On top of these 256 dimensions, the scaled intensities were added as an additional dimension. The first peak was set to intensity 2.0 and the encoded precursor ion mass. The spectral sequences were padded to a maximum length of 501, resulting in a final matrix with a shape of 501x257.

SMILES were encoded with the cddd package (Winter et al., 2019), which is based on a pretrained continuous chemical descriptor space. The number of 60 different functional groups was extracted using prebuilt rdkit functions and the number of sugars was identified by sugar removal utility (Schaub et al., 2020) with the command *-t “3” -remTerm “false”*. Atom counts from molecular formulas were extracted with the molmass package. These numbers were then scaled to floating numbers to encode the information for the neural network.

The neural network architecture (**Figure 1**) has a total of 33 million parameters and is based on 5 standard transformer encoder layers (16 heads and 2048 units for feed forward) as described in Vaswani et al., (2017) which are feeding into a temporal convolutional neural network (TCN) (Bai et al., 2018) with a receptive field of 883, a kernel size of 8 and 256 filters. This is followed by 2x2 dense layers that produce two outputs of shape 512 and 71. The architecture is implemented in tensorflow (version 2.11.0, Abadi et al., 2015) and the TCN is implemented by the keras-tcn package (Bai et al., 2018). The training was performed on one Nvidia Tesla V100S GPU with 32 Gb RAM on the IBMP computing cluster. Training progress was logged with the package wandb (version 0.13.5). A hyperparameter search was performed with keras-tuner package (Chollet and others, 2015) in the random search mode for 99 trials, with one execution *per* trial and 4 epochs (see also **Data and code availability**). In addition, manual inspections were performed to find optimal parameters. The final training was stopped after 50 epochs, as the model performance did not significantly improve with longer training (**Figure S1**).

**Figure 1.**
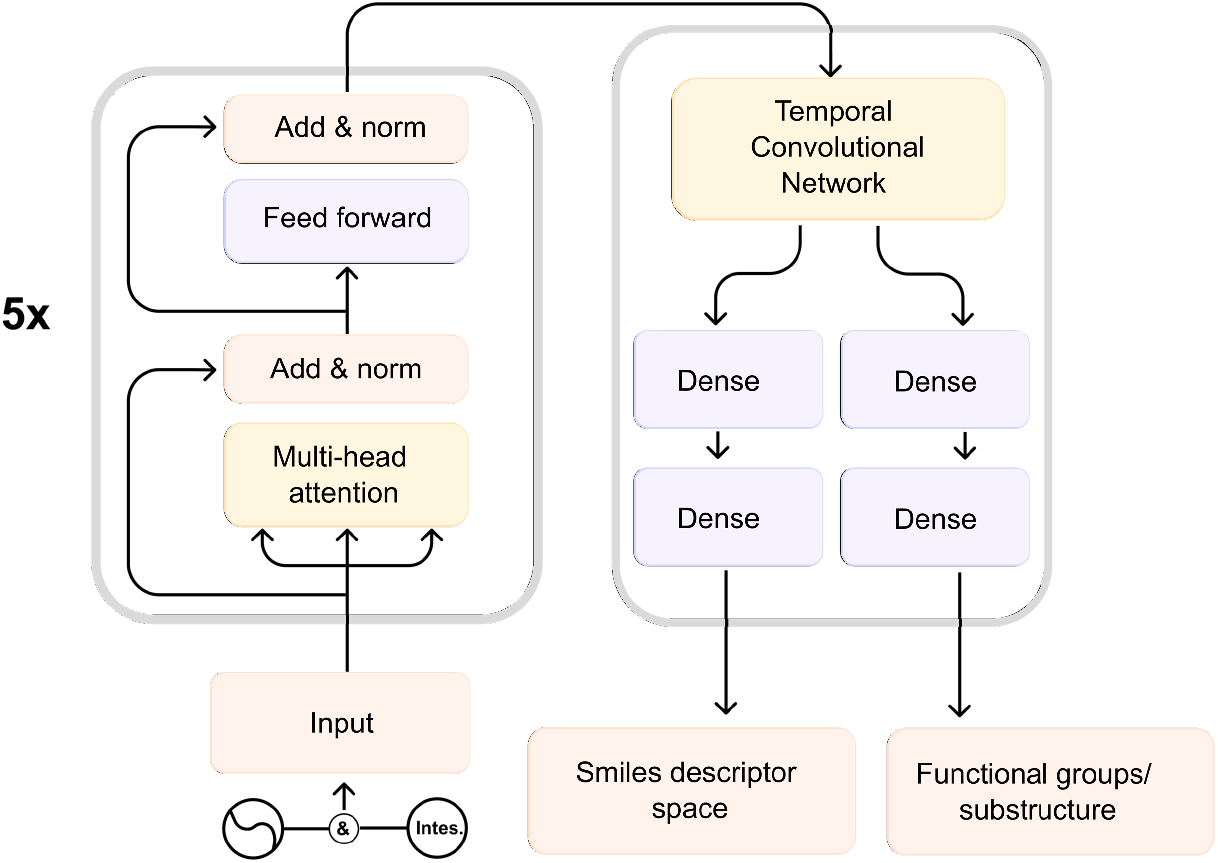
General architecture of Mass2SMILES. Encoded spectra are fed into the five transformer encoder layers and then processed as part of a temporal convolutional neural network. The final two outputs are produced by dense layers, which returns the chemical descriptor space encoding for the SMILES strings and the encoded 60 functional groups as well as number of atoms and the adduct type.

Network analysis was performed with the matchms (version 0.15.0) and matchms extras (version 0.4.1) using the modified cosine score (tolerance=0.01) as similarity measure. Molecular networks were created (score_cutoff=0.7, max_links=10), exported to cytoscape and Mass2SMILES annotations were visualized by the chemVIz2 plugin.

## Results and Discussion

Training of the final model of Mass2SMILES was accomplished within one day and two hours for 83,358 spectra on one Nvidia Tesla V100S GPU. The model loss was evaluated by the calculation of the Mean Absolute Error (MAE) and of the Mean Squared Error (MSE). The final loss MAE for SMILES descriptor space was of 0.18 (**Figure 2A**) and the MSE was of 0.06, whereas the MAE calculated for the functional groups was of 0.004 and the MSE of 0.00006 (**Figure 2D**). The final loss that was achieved on the validation data-set for the SMILES descriptor space was of 0.24 (**Figure 2B**) and the MSE was of 0.1, whereas for the functional groups MAE was of 0.004 and MSE of 0.0001 (**Figure 2C**). The final model was inferred on CPU through a docker container with the command: *docker run -v c:/Users/delser/mass2smiles/:/app mass2smiles:transformer_v1 conda run -n tf python app/mass2smiles_transformer.py input_file.mgf/app*, which on average takes two seconds processing time for one pair of structure and functional groups on our machine. This makes it suitable to predict chemical structures for large-scale metabolomics studies, using GPU for inference could even drastically increase inference speed. The predictions from the 236 positive ion mode spectra of the CASMI 2022 contest retrieved 2 true structures **(Table 2**) and one candidate with all true numbers of functional groups (**Table 3**) (**Data and code availability: Supplemental Data S1)**. Interestingly, the number of correctly predicted functional groups was not necessarily reflected in the *Tanimoto* similarity. Training two different models for each output did not improve accuracy but rather slightly reduced performance e.g., from two true CASMI 2022 predictions to zero (the best one having a 0.96 *Tanimoto* similarity to the true structure and an average *Tanimoto* similarity of 0.38). The average *Tanimoto* similarity achieved from the final model (with 2 outputs) on this dataset was of 0.4, with two structures having a similarity higher than 0.9 and 10 more than 0.7. On average there were 51 true numbers of functional groups out of 60. The molecular formula estimation resulted into 8 true hits, whereas when the predicted SMILES were converted into molecular formulas, 7 true hits were retrieved. For 59 out of 236 molecules, the network was able to predict the true number of heteroatoms, which on average resulted into 7 out of 8 possible heteroatom numbers.

**Figure 2.**
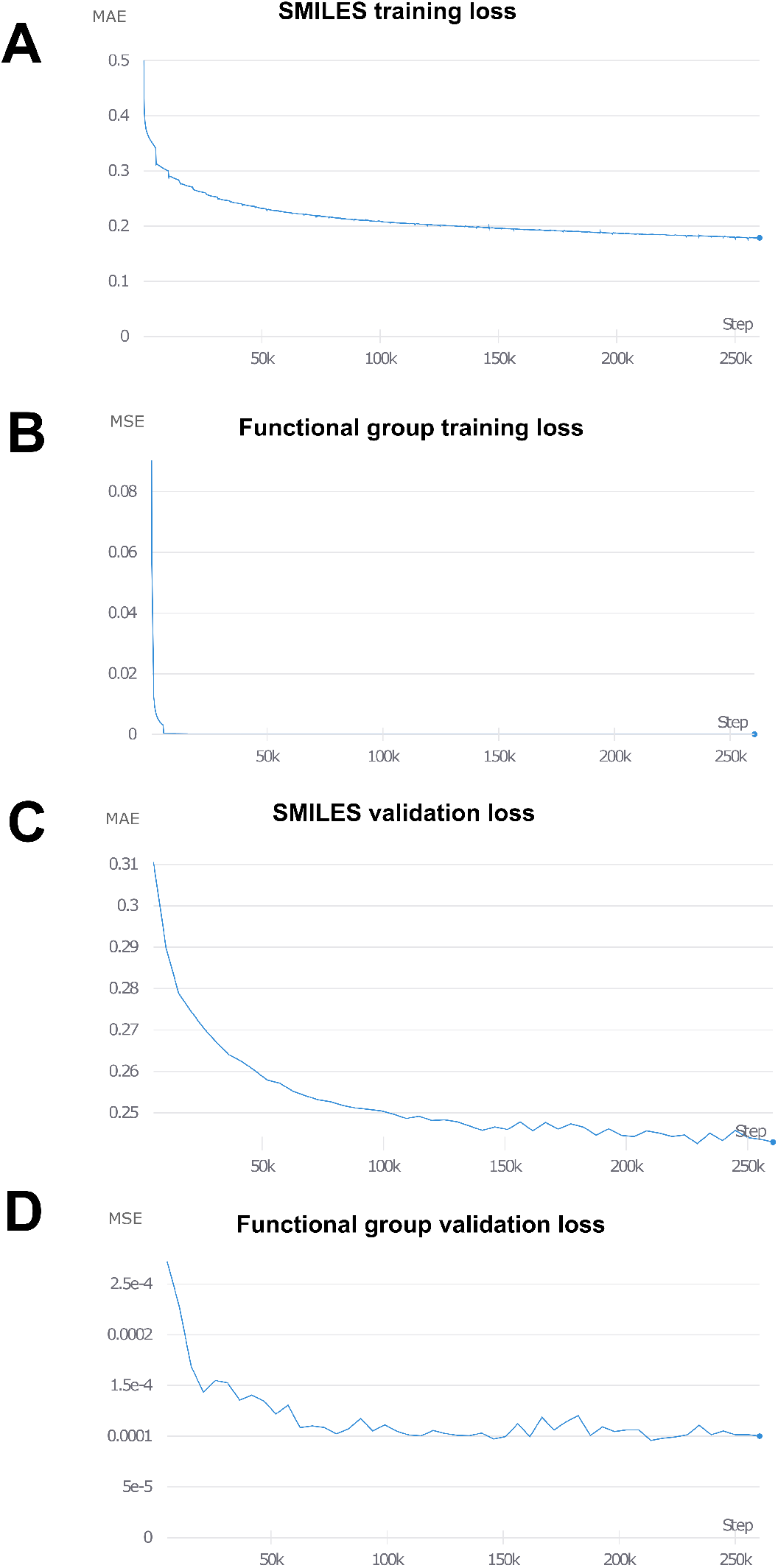
The training progress on the final model as tracked by wandb. The mean absolute error (MAE) and the mean squared error (MSE) are shown according to the number of training steps. The training was stopped after 50 epochs, as further training did not seem to improve the performance, one epoch is comprised of 5210 steps.

**Table 2.**
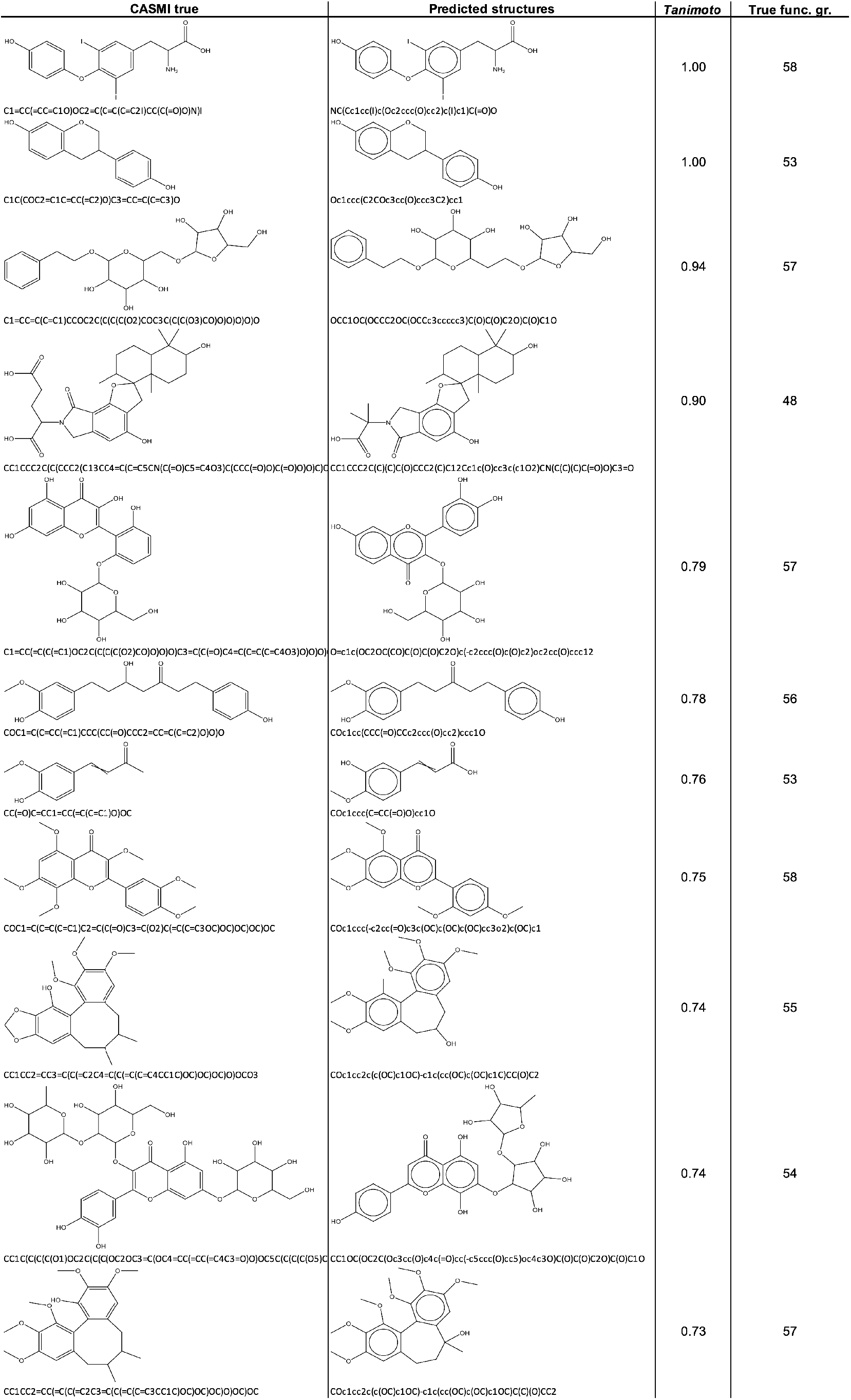
The top predictions on the CASMI 2022 positive mode dataset sorted by *Tanimoto* similarity.

**Table 3.**
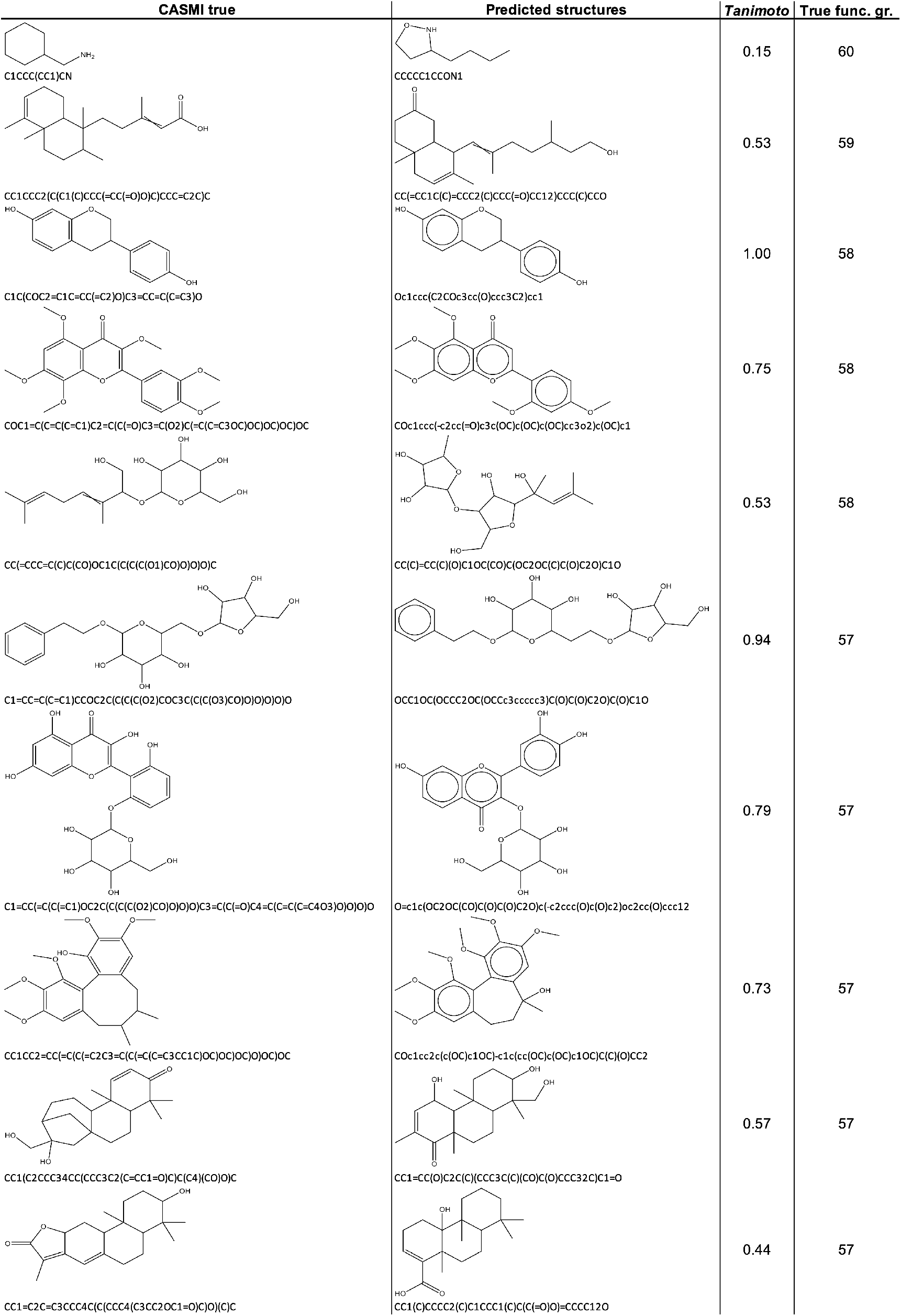
The top predictions on the CASMI 2022 positive mode dataset sorted by the number of true functional groups. The true number of functional groups does not necessarily align with *Tanimoto* similarity.

From the 744 validation spectra, Mass2SMILES was able to predict 7 true structures (**Table 4**), followed by 14 predictions with *Tanimoto* scores above 0.9, 62 with *Tanimoto* scores above 0.7. The average *Tanimoto* similarity between predicted and true structures was of 0.39. Moreover, the model was able to correctly predict the exact presence of functional groups for 17 spectra (**Table 5**), whereas on average the model predicted 54 true numbers of across the dataset (out of a maximum of 60) (**Data and code availability: Supplemental Data S2)**. Interestingly, the model was able to correctly predict 436 adducts, for 11 molecules it found the true adduct and the molecular formula. For 22 molecules, it found the true molecular formula alone and for 187 the true number of heteroatoms. When converting the predicted SMILES into molecular formulas it retrieved 63 true hits out of 744.

**Table 4.**
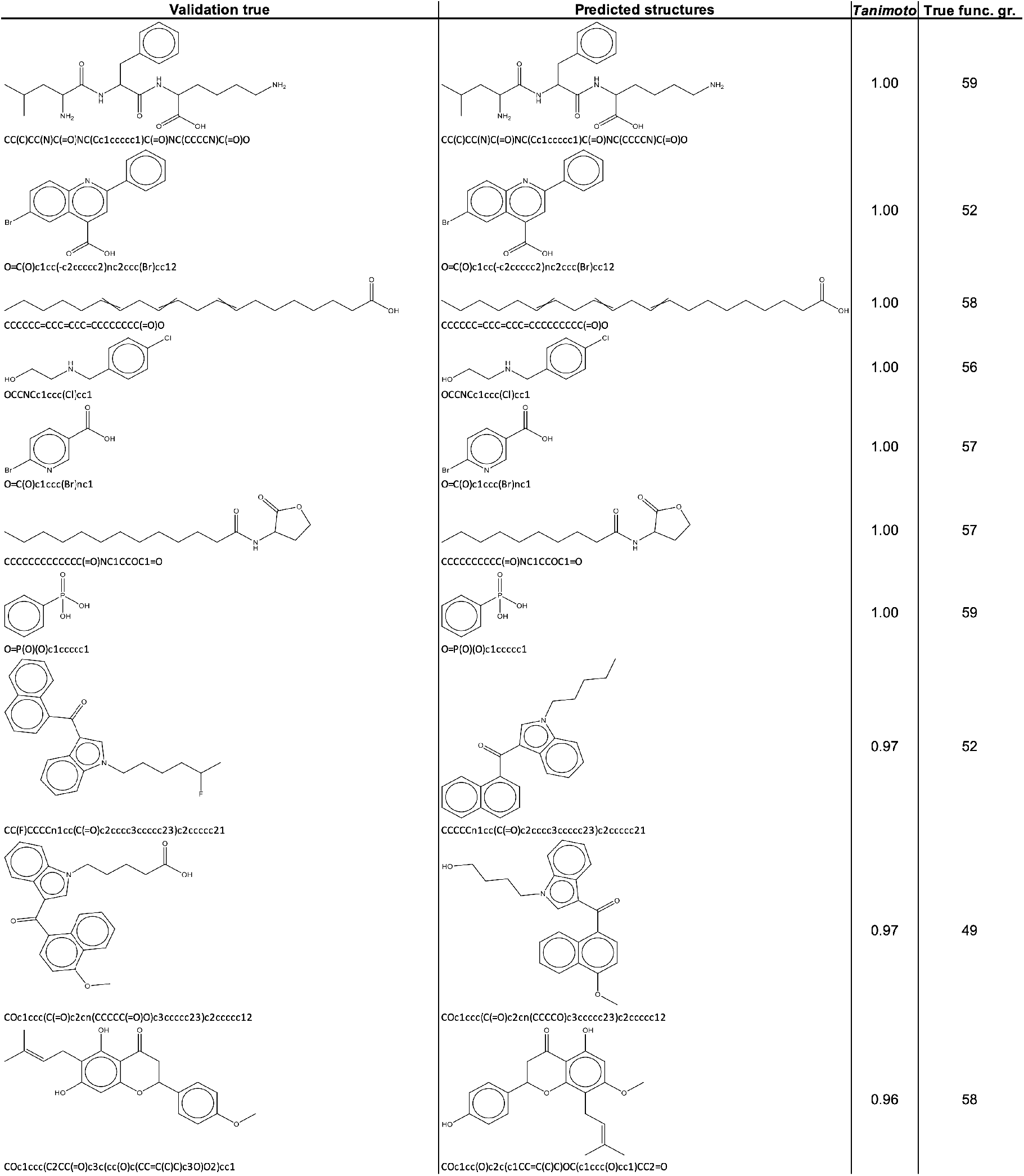
The top predictions on the validation dataset sorted by *Tanimoto* similarity.

**Table 5.**
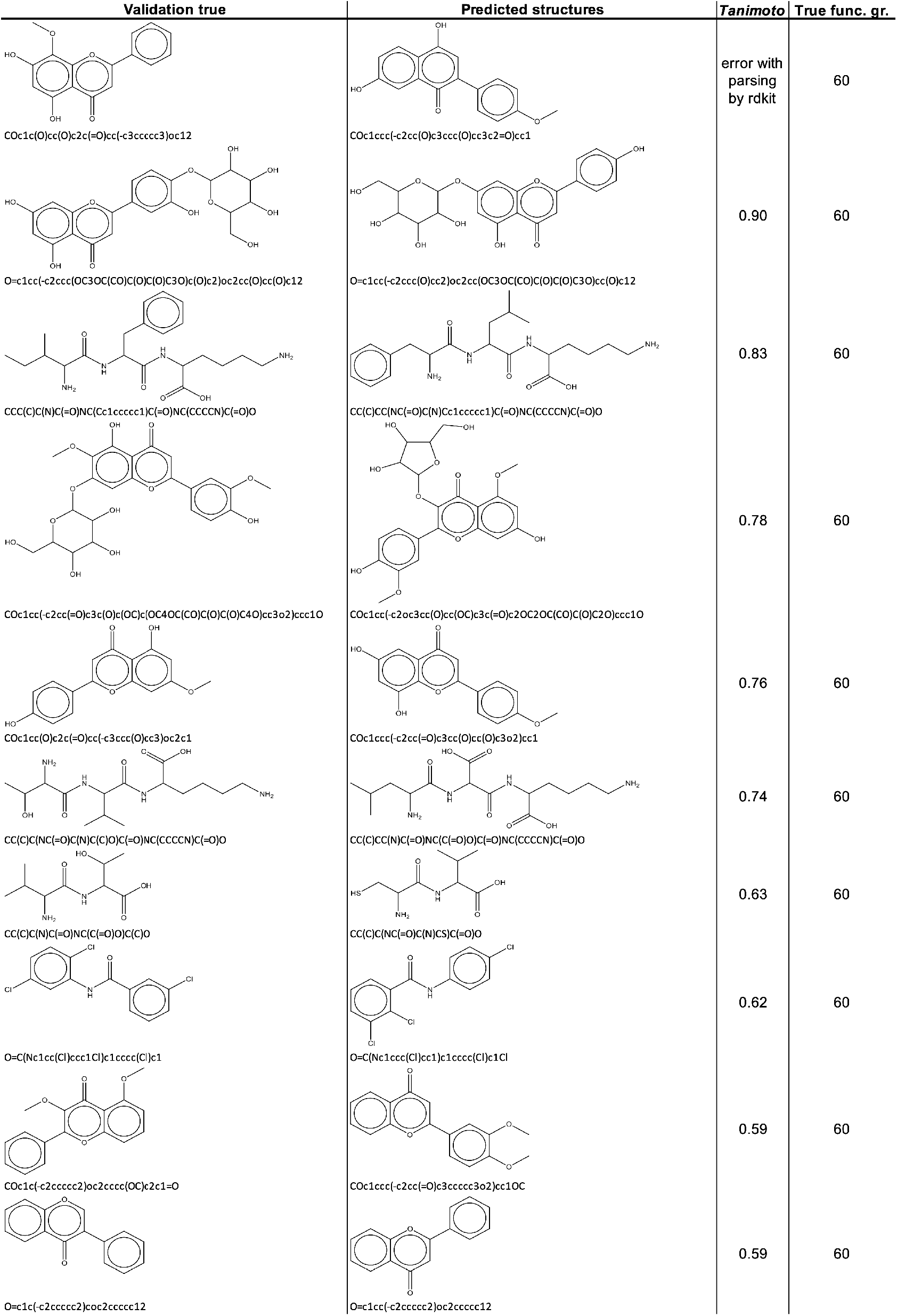
The top prediction on the validation dataset, sorted by the number of true functional groups. The true number of functional groups does not necessarily align with *Tanimoto* similarity.

We then also examined Mass2SMILES on a metabolomics dataset acquired for 20 *Nicotiana* species (Elser et al., 2022) to better judge of its performance on a real use case study. For this, we first predicted all the structures across the whole dataset. On a Intel Xeon E5-2630 v2 @ 2.6 GHz CPU, this took 9 hours and resulted in 16616 smiles out of 17902 spectra, most likely the spectra without predictions contained less than 6 peaks and were therefore automatically discarded by Mass2SMILES. In general, the model produces valid SMILES in most of the cases e.g. for the 744 validation spectra only 10 created errors with parsing by rdkit. For 457 features on the *Nicotiana* dataset, the molecular formula was identical with the one predicted by SIRIUS which is frequently is described as a gold standard method to perform this task. These features were then selected to predict the ClassyFire classes (Djoumbou Feunang et al., 2016) which were then compared to the ones predicted in parallel by CANOPUS (Dührkop et al., 2021) directly from the MS/MS spectra. For 235 of these, the superclass was identical and for 176 the class prediction matched. When inspected with in further details, classes that had a mismatch were frequently very close and in a lot of cases the Mass2SMILES prediction was even closer to the actual true structure (**Table 6**). Interestingly, Mass2SMILES was able to annotate structures that did not retrieve database hits, even for very common molecules such as nicotine (**Table 6**). This shows that neural networks such as implemented as part of Mass2SMILES could possibly provide alternative solutions to computationally intensive database searches. We observed for several spectra very accurate annotations that did not yield any database hits but were predicted to belong to these classes e.g. terpenoids, coumarins, flavonoids, O-acyl-glucoses or O-acyl-glycerols (**Data and code availability: Supplemental Data S3** and **Table 6**). Some molecules did not yield a CANOPUS annotation but could nonetheless be accurately predicted with Mass2SMILES (**Table 6**).

**Table 6.**
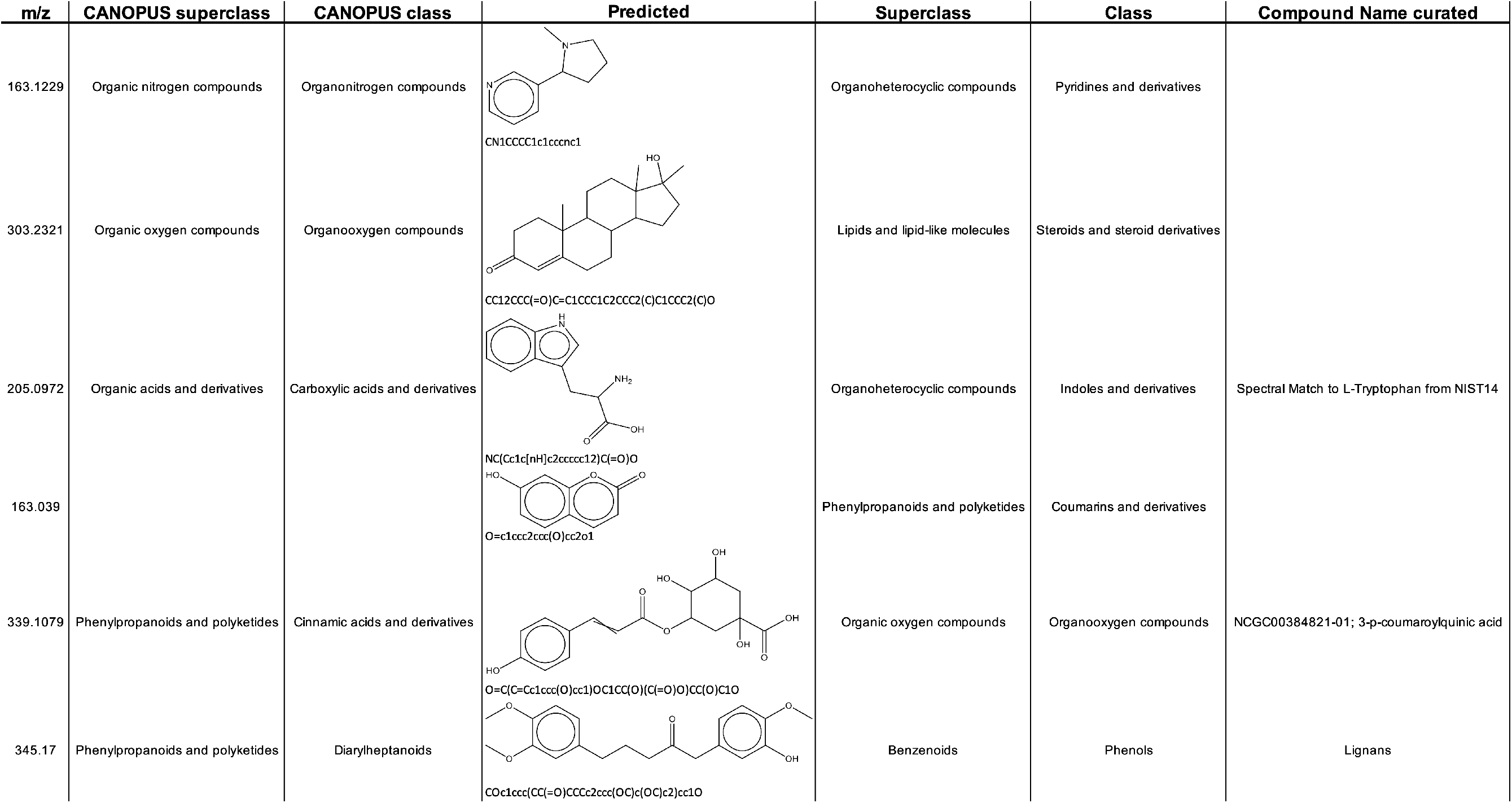
Selected predictions obtained for the metabolomics dataset on *Nicotiana* species (Elser et al., 2022). First, predicted SMILES were converted into molecular formulas and then compared with the predictions from SIRIUS, if consistent, SMILES were further converted with ClassyFire into chemical classes. Some examples of class predictions that did not match with CANOPUS ones are additionally depicted. If blank, no annotation was generated in the study from Elser et al. (2022).

As an additional case study, we processed metabolomics data and predicted structures with Mass2 SMILES for MS/MS spectra collected from a dataset of 9 bryophyte analyzed and reported earlier by Peters et al. (2018). This dataset had further been previously used to test the performance of the MSNovelist *de novo* structure elucidation tool (Stravs et al., 2022). **Figure 3** shows a molecular network that comprises the MS/MS feature 377 for which a flavonoid-like structure had been initially predicted as part of the study reporting the performance of MSNovelist (Stravs et al., 2022). It is striking to see that several flavonoid-related structures were predicted for this network, by Mass2SMILES. The structure, the number of aromatic hydroxyl groups (Mass2SMILES: 3/ MSNovelist: 5) and the number of benzene rings (Mass2SMILES: 2/ MSNovelist: 3) predicted by MSNovelist does not correspond with the ones predicted by Mass2SMILES for this feature 377. Nonetheless, both Mass2SMILES and MSNovelist converge on a flavonoid like structure for this feature. Molecular networks with structures as depicted in **Figure 3** may hence be combined in future studies with Mass2SMILES or MSNovelist predictions to further assist in the annotation of unknowns as well as to give insights into the compound class and dominating functional groups.

**Figure 3.**
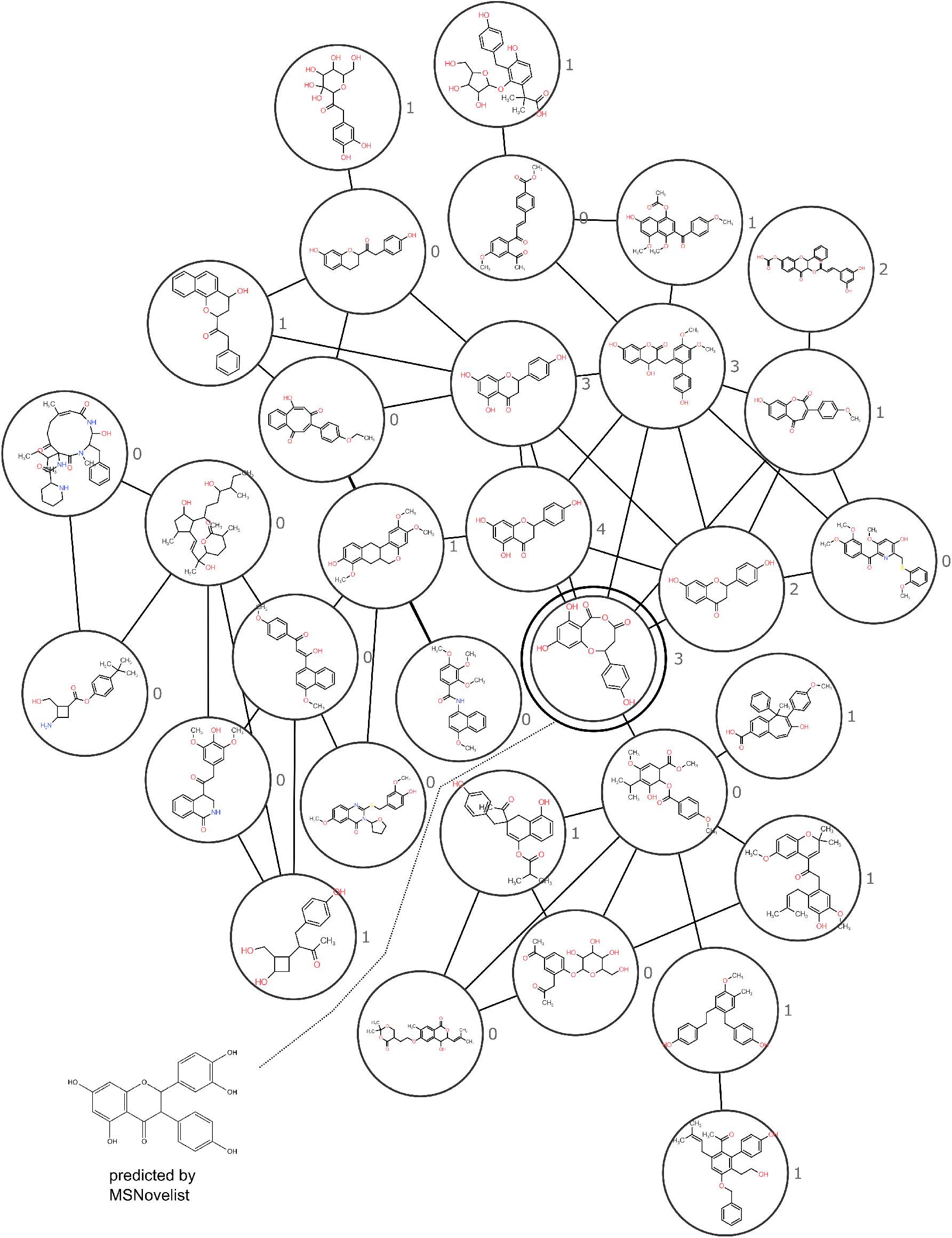
A flavonoid related molecular network from bryophytes, annotated with Mass2SMILES. This network was constructed on metabolomics data by Peters et al. (2018). The same dataset had been previously used for structure prediction as part of the publication of MSNovelist (Feature 377, Stravs et al., 2022). Numbers indicate the number of predicted aromatic hydroxyl groups, this number is not in line with the structure predicted by MSNovelist (bottom left). The double circled structure (Feature 377), is the proposed structure by Mass2SMILES.

One drawback of Mass2SMILES is that it relies on the cddd package (Winter et al., 2019) which runs on relatively old Python (3.6) and tensorflow (1.10) versions, future models should be built on a pretrained SMILES transformer model such as ChemBERTa-2 (Ahmad et al., 2022) or a transformer model that has been trained on a random to canonical SMILES translation task such as is the cddd model, but with recurrent neural networks. The major limitation we see to further improve the accuracy of Mass2SMILES, however, is the lack of comprehensive publicly available annotated MS/MS data to better cover the extremely structurally diverse chemical space of natural products. We expect that including high quality MS/MS data from databases such as METLIN or Mzcloud would greatly increase the performance of the overall model. In addition, future progress in molecular dynamics calculations such as QCxMS (Koopman and Grimme, 2021) and increased computing power would offer a new potential for creating more sophisticated models. With the increasing availability of large-scale annotated MS/MS data, the use of large language models such as LLaMA (Touvron et al., 2023), GPT-NeoX (Black et al., 2022) or Chinchilla (Hoffmann et al., 2022) seems to be a highly promising methodological avenue to train a new generation of structure prediction models in the future.

## Conclusions

Mass2SMILES is a novel deep learning-based approach for the annotation of MS/MS spectra with SMILES, which in addition also predicts the number of several functional groups present in a molecule. It is also able to predict the adduct type and gives an estimation of the molecular formula. This software can easily be applied to large metabolomics datasets and may represent an alternative to computationally intensive database searches. We demonstrate the capabilities of Mass2SMILES on the CASMI 2022 dataset, as well as on a previously reported large scale metabolomics dataset. We expect that this tool will aid the metabolomics community in further illuminating the large amount of dark matter present in current experiments.

## Acknowledgments

The authors would like to acknowledge the High-Performance Computing Center of the University of Strasbourg for supporting this work by providing scientific support and access to computing resources, notably funded by the Equipex Equip@Meso project (Programme Investissements d’Avenir) and the CPER Alsacalcul/Big Data. In addition, we would like to acknowledge access to computing resources at the Institute of Molecular Biology of Plants (IBMP), CNRS | University of Strasbourg (IBMP). **Funding:** D.E., and E.G. were funded by the CNRS. D.E. and E.G. were supported by a IdEx (Investissement d’Avenir) grants from the University of Strasbourg, with a IdEX PhD fellowship to D.E and a IdEx Grant Recherche Exploratoire to E.G.

## Author contributions

D.E. conceived the study, performed coding, analyzed the data. F.H. and E.G. supervised the study. All authors equally contributed to writing the manuscript.

## Data and code availability

Supplemental Data and the Docker container are available on Zenodo https://doi.org/10.5281/zenodo.7883491 All scripts used in this study are available at the Github repository: https://github.com/volvox292/mass2smiles. All data needed to evaluate the conclusions in the paper are further present in the paper and/or the Supplementary Materials.

---

**Figure S1.**
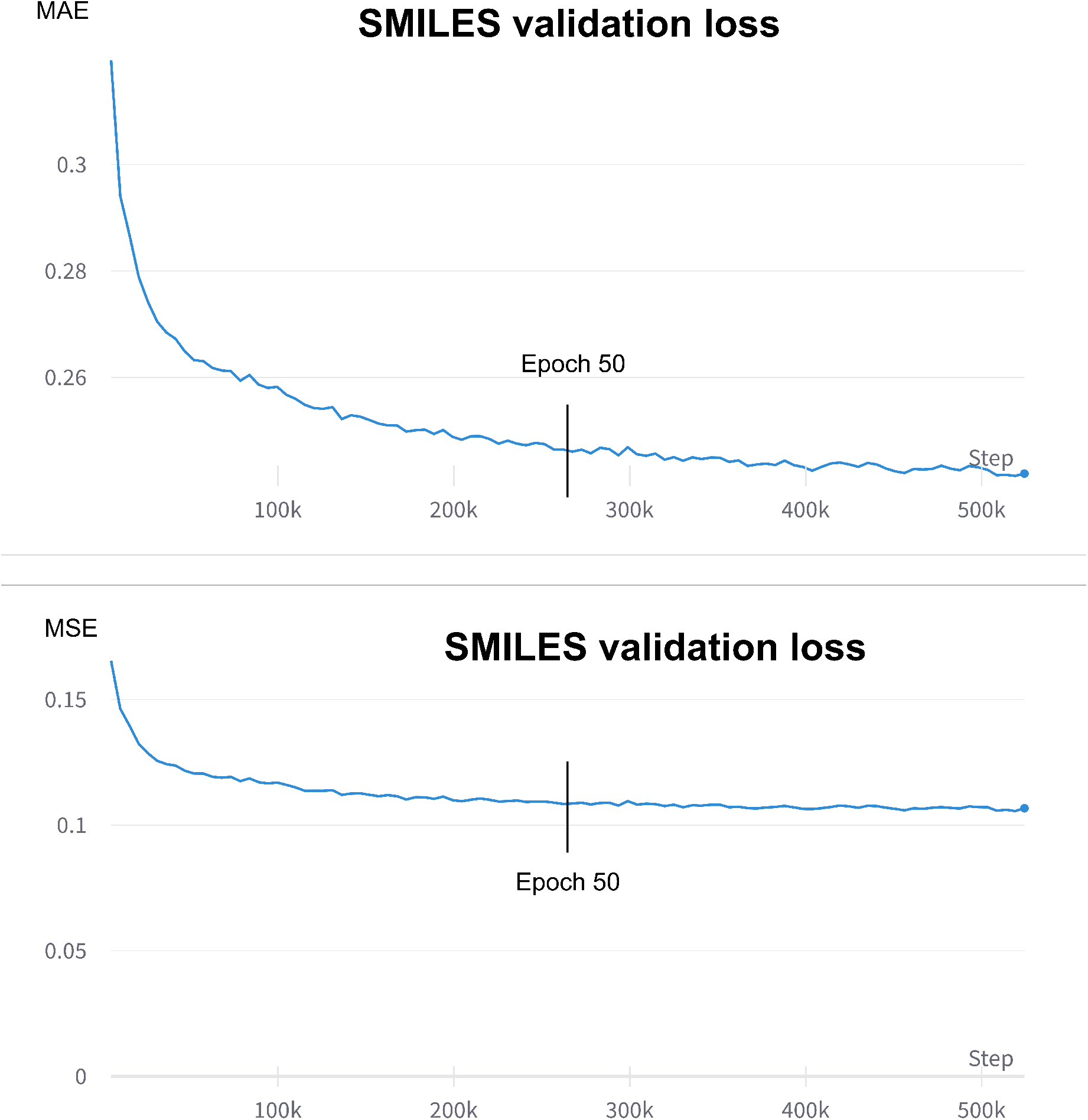
Example of a Mass2SMILES training run with 100 epochs. Longer training did not significantly improve the overall model performance. The duration of 50 epochs was therefore chosen as good compromise between model performance and training duration.

